# The genetic architecture and genomic context of glyphosate resistance in *Amaranthus tuberculatus*

**DOI:** 10.1101/2020.08.19.257972

**Authors:** J.M. Kreiner, P.J. Tranel, D. Weigel, J.R. Stinchcombe, S.I. Wright

## Abstract

Much of what we know about the genetic basis of herbicide resistance has come from detailed investigations of monogenic adaptation at known target-sites, despite the increasingly recognized importance of polygenic resistance. Little work has been done to characterize the broader genomic basis of herbicide resistance, including the number and distribution of genes involved, their effect sizes, allele frequencies, and signatures of selection. In this work, we implemented genome-wide association (GWA) and population genomic approaches to examine the genetic architecture of glyphosate resistance in the problematic agricultural weed, *Amaranthus tuberculatus*. A GWA was able to correctly identify the gene targeted by glyphosate, but when we statistically controlled for two target-site genetic mechanisms, we found an additional 250 genes across all 16 chromosomes associated with non-target site resistance (NTSR). The encoded proteins had functions that have been linked to non-target site resistance (NTSR), the most significant of which is response to chemicals, but also showed pleiotropic roles in reproduction and growth. The architecture of NTSR was enriched for large effect sizes and low allele frequencies, suggesting the role of pleiotropic constraints on its evolution. The enrichment of rare alleles also suggested that the genetic architecture of NTSR may be population-specific and heterogeneous across the range. Despite their rarity, we found signals of recent positive selection on NTSR-alleles by both window- and haplotype-based statistics, and an enrichment of amino-acid changing variants. In our samples, genome-wide SNPs explain a comparable amount of the total variation in glyphosate resistance to monogenic mechanisms, even in a collection of individuals where 80% of resistant individuals have large-effect TSR mutations, indicating an underappreciated polygenic contribution to the evolution of herbicide resistance in weed populations.

## Introduction

In theory, it should be easy to understand the genetic basis of herbicide resistance in weeds, because herbicides typically intentionally target a known biochemical pathway, and specific gene products. In practice, however, herbicide resistance in evolved weed populations is complex. Agricultural weed populations evolve resistance not only through well characterized large-effect mutations that alter the interaction of the herbicide with the target enzyme (target-site resistance, TSR), but also through other mutations across the genome (non-target-site resistance, NTSR), often with smaller individual effects (Délye, 2013; Kreiner, Stinchcombe, & Wright, 2018; Tranel, Riggins, Bell, & Hager, 2011). While much work has been done to elucidate causative TSR mutations and their frequency in experimental and agricultural weed populations (Heap, 2010), we are only beginning to learn about the number of loci involved and the associated distribution of allelic effect sizes, frequencies, and the fraction of phenotypic variance explained by NTSR (Van Etten, Lee, Chang, & Baucom, 2019). Here we combine genome-wide association approaches with population genomics to dissect the genetic architecture and genomic context of glyphosate resistance.

Efforts to better characterize the genetic basis of NTSR will help discover genetic markers for managing the spread of resistance. With a catalogue of genome-wide allelic effects on resistance, GWA methods can inform prediction of how populations, based on their genetics, may respond to selection from future herbicide pressures. More broadly, uncovering the genetics of NTSR will contribute to our understanding of the relative importance and consistency of small- and large-effect alleles to rapid adaptation, while also shedding light on how strong selection shapes genome-wide variation of natural weed populations.

NTSR can be broadly defined as all alleles that provide tolerance or resistance to herbicides unexplained by and unlinked to TSR, regardless of mechanism—by providing direct protective effects, compensentating for herbicide toxicity through downstream pathways, or mediating the costs and/or benefits of TSR alleles—and we use this conceptual approach in this work. It is likely that such mechanisms are similarly important as TSR in conferring herbicide resistance across agriculturally important ranges (e.g. Delye, Gardin, Boucansaud, Chauvel, & Petit, 2011), as several studies have looked for causative mutations in the gene encoding the herbicide-targeted protein in resistant populations, but failed to find one (Délye, 2013; Guo et al., 2015; Van Etten et al., 2019; Van Horn et al., 2018). In contrast to a single large-effect TSR locus, where resistance alleles are often assumed to have arisen *de novo*, the polygenic basis of resistance with many small-effect loci is more likely to draw from genome-wide standing genetic variation (Kreiner et al., 2018; Paul Neve, Vila-Aiub, & Roux, 2009). Thus, NTSR mechanisms may even allow for naive populations to have some level of standing variation for resistance to herbicides. NTSR is a challenge for management not only because of its mysterious genetic basis and possible presence in naive populations, but also due to pleiotropic effects of resistance alleles conferring cross-resistance to multiple herbicide modes (Preston, 2004; Preston, Tardif, Christopher, & Powles, 1996; Yu, Abdallah, Han, Owen, & Powles, 2009; Yu, Cairns, & Powles, 2007). With increased incidence of cross-resistance within individuals and populations (Heap, 2010), combined with reactive management through applications of tank mixes of different classes of herbicides, NTSR is likely to only increase in importance and prevalence.

A handful of gene families—encoding enzymes such as cytochrome P450s, monooxygenases, glycosyltransferases, and glutathione S-transferases, or ABC transporters—have been repeatedly implicated in conferring NTSR, especially through the action of differential gene expression (Yuan, Tranel, & Stewart, 2007). These NTSR genes are thought to act by altering herbicide penetration, translocation, accumulation at the target site, as well as through offering protection from herbicide effects (Délye, 2013; Moretti et al., 2018), the function of several of which have been experimentally validated (Ito & Gray, 2006; Xi, Xu, & Xiang, 2012). However, the genomic, rather than transcriptomic, basis of NTSR is unknown in most species and for most classes of herbicides (but see Cummins et al., 2013; Van Etten et al., 2019).

While increasingly the genetic repeatability of herbicide resistance is being investigated with genomic approaches (Flood et al., 2016; Kreiner et al., 2019; Küpper et al., 2018; Leslie & Baucom, 2014; Molin, Wright, Lawton-Rauh, & Saski, 2017; Van Etten et al., 2019), traditional genome-wide association (GWA) approaches have yet to be applied to explicitly characterize the genetic architecture of herbicide resistance, and the repeatability of alleles implicated across populations (but for related approaches, see Benevenuto et al., 2019; Van Etten et al., 2019). In other systems, GWAS have been widely and successfully used to identify new candidate genes, characterize the architecture of key traits, make predictions about phenotypes in studied and novel populations, and inform breeding strategies through genomic selection (e.g. Bock, Kantar, Caseys, Matthey-Doret, & Rieseberg, 2018; Epstein et al., 2018; Exposito-Alonso et al., 2019, 2018; Spindel et al., 2015; Swarts et al., 2017).

Glyphosate-based herbicides, commonly referred to by the brand name Round-up™, are widely used across North America to suppress weed populations, especially in soy and corn fields. Glyphosate targets a step in the synthesis of aromatic amino acids, carried out by 5-enolpyruvylshikimate-3-phosphate synthase (EPSPS). These aromatic amino acids are required not just for protein synthesis, but for the synthesis of many natural compounds such as hormones, pigments, and cell wall components, and their production draws on more than 30% of photosynthetically fixed carbon (Maeda & Dudareva, 2012). Glyphosate herbicides were commercialized in the 1970s with widespread deployment of glyphosate-resistant crops nearly 20 years later (W. G. Johnson, Davis, Kruger, & Weller, 2009). Since then, the incidence of glyphosate resistance has increased exponentially, and is third to only ALS-inhibiting and Photosystem II-inhibiting herbicides in the number of resistant species (Heap, 2010). Meanwhile, glyphosate is still the most widely used herbicide across North America—due to low operational costs and glyphosate-resistant cropping systems—despite ubiquitous resistance in weed populations, especially in the genus *Amaranthus* (Duke & Powles, 2008; Vencill, Grey, Culpepper, Gaines, & Westra, 2008).

*Amaranthus tuberculatus* has been investigated by weed scientists for decades due to its problematic nature in agricultural fields and increasing reports of multiple-resistance (Bell, Hager, & Tranel, 2013; Bernards, Crespo, Kruger, Gaussoin, & Tranel, 2012; Matthew J. Foes, Tranel, Loyd M. Wax, & Edward W. Stoller, 1998; Patzoldt & Tranel, 2007; Patzoldt, Tranel, & Hager, 2002, 2005; Shergill, Bish, Jugulam, & Bradley, 2018; Tranel et al., 2011). More recently, a variety of genomic resources (a high quality reference genome, large resequencing dataset, phenotyping for glyphosate resistance) have been assembled (Kreiner et al., 2019), building on earlier efforts (Lee et al., 2009; Riggins, Peng, Stewart, & Tranel, 2010). Recently, we characterized population structure, demographic history, and signals of monogenic selection at EPSPS and its related amplification in response to glyphosate, across natural and agricultural populations (Kreiner et al., 2019). Our previous work was a targeted investigation of repeated origins of monogenic target-site resistance, particularly signals of positive selection from glyphosates around the EPSPS gene in individuals with high EPSPS copy number but did not attempt to characterize polygenic signals of selection for herbicide resistance. With TSR mechanisms incompletely explaining phenotypic resistance levels in those samples (Kreiner et al., 2019), and reports of non-target site resistance to glyphosate in the species (Nandula, Ray, Ribeiro, Pan, & Reddy, 2013), in the current study we sought to identify genome-wide effects of resistance and signals of adaptation to herbicides.

We implemented GWA and population genomic approaches for characterizing the genetic architecture of glyphosate resistance in *Amaranthus tuberculatus*. Specifically, we aimed not only to identify putative non-target site loci across the genome, but also to explore the relative importance of this polygenic architecture compared to previously characterized target-site resistance mutations in our dataset. Furthermore, we sought to understand modes of selection on polygenic resistance. As expected, a GWA for resistance to glyphosate in *Amaranthus tuberculatus* without TSR covariates shows the largest genome-wide peak on chromosome 5 near the known-target, EPSPS. To identify putative NTSR loci, we included both of the known TSR mechanisms as covariates in the GWAS (as in Atwell et al., 2010; Sasaki, Köcher, Filiault, & Nordborg, n.d.); this revealed hundreds of other alleles with genome-wide significance after correcting for multiple testing (FDR) and relatedness among individuals. Furthermore, we compared our observations to an empirical null that allowed us to evaluate the extent to which differences in the genetic architecture of resistance across populations may be confounded with population structure. Against this null expectation, we asked whether the architecture we identified was enriched for relevant NTSR-related functions and population genomic signals of selection, and evaluated its importance against widespread TSR mechanisms. This work contributes to our understanding of the heterogeneity in the genetic architecture of resistance across populations, its importance compared to TSR mechanisms, and selective processes that have governed its evolution.

## Materials & Methods

### Amaranthus tuberculatus resequencing & phenotype data

Resequencing and phenotype data were obtained from a published study (Kreiner et al., 2019), which included a high-quality female reference genome for the species. Whole-genome Illumina sequencing data are available at European Nucleotide Archive (ENA) (project no. PRJEB31711) (Kreiner, Giacomini, Bemm, Waithaka, Regalado, Lanz, Hildebrandt, Sikkema, Tranel, Weigel, Stinchcombe, Wright, 2019b), and the reference genome and its annotation available at CoGe (reference ID = 54057) (Kreiner, Giacomini, Bemm, Waithaka, Regalado, Lanz, Hildebrandt, Sikkema, Tranel, Weigel, Stinchcombe, Wright, 2019a). There were 158 agricultural genomic samples, collected from 8 fields with high *A. tuberculatus* densities across Missouri and Illinois in the Midwest United States, and from counties with newly resistant populations in Ontario, Canada, Walpole Island and Essex. Lastly, 10 samples from naturally occurring, non-agricultural Ontario populations were included, totaling 168 resequenced samples.

As described in Kreiner et al., 2019, phenotyping for glyphosate resistance was performed from 3-5 offspring individuals per line of field-collected specimens. Three weeks after a glyphosate application of 1,260 g glyphosate (WeatherMax 4.5 L; Monsanto) per hectare, plants were rated for resistance on a scale of 1 to 5 based on a visual scale of percent injury, with 1 indicating complete plant necrosis and 5 indicating no apparent damage. A subset of these phenotyped offspring individuals were then selected for resequencing, with the goal of representing both susceptible and resistant individuals across populations (i.e., the distribution of resistance within our set of sequenced individuals was not chosen to reflect population frequencies). Individual-level resistance ratings were directly used in the GWAS, as opposed to family-level means, as variance in resistance within a family may be encompassed by differences at many loci genome-wide. We analyzed GWAs with resistance using these discrete 1-5 ratings.

### Genetic TSR characterization & Multiple Regression

We previously characterized TSR mechanisms in (Kreiner et al., 2019). Briefly, after SNP calling and filtering, we located the known TSR mutation codon *EPSPS* Pro106Ser by submitting a validated EPSPS complete mRNA coding sequence (genbank sequence: FJ869881.1) to a BLAST (Altschul, Gish, Miller, Myers, & Lipman, 1990) search to our reference genome. Individuals containing the Pro106Ser mutation were all heterozygous, and thus the *EPSPS* Pro106Ser covariate was coded based on presence/absence (0/1). Individual EPSPS copy number was estimated by dividing the median coverage at every base-pair position within the EPSPS gene, by the median genome-wide coverage (since the median coverage should be a good proxy for coverage of single copy genes genome-wide) and taking the gene-wide average.

For estimating the proportion of phenotypic variance in resistance explained (PVE) by TSR mechanisms, we used a simple linear regression of resistance on Pro106Ser presence/absence and *EPSPS* copy number. We decided not to include the kinship matrix as a covariate in this regression, as validated large-effect TSR mutations are unlikely to be driven by population structure and our aim was to avoid attributing variance in resistance explained by TSR to genome-wide relatedness. This should lead to a relatively conservative estimate of the importance of NTSR mechanisms compared to TSR mechanisms. While ideally, we would be able to extract this information directly from the GWA of NTSR that includes both genetic covariates, it is infeasible to do so given current GWA software packages, which often have the goal of controlling for covariates rather than estimating the variance explained as an object of study.

To estimate the SNP heritability of glyphosate resistance, we additionally used the R package “lme4qtl” to run a multiple regression of resistance onto both TSR genetic mechanisms including the kinship matrix used in our GWA as a random-effect covariate, using the function “relmatLmer”.

### Linear mixed GWA models & preprocessing & GO enrichment

Filtered VCFs were obtained from (Kreiner et al., 2019). Briefly, freebayes-called SNPs were filtered based on missing data (<=20% missing), repeat content, allelic bias (<0.25 and >0.75), read paired status, and mapping quality (> Q30). Six individuals were removed due to excess missing data, leaving 162 for baseline analyses. Further investigations into identity-by-descent in these 162 samples showed some individuals with particularly high levels of relatedness, some of which was unaccounted for even when including a relatedness matrix in genome-wide association tests. We thus removed another 7 individuals, resulting in a final sample size of 155.

We implemented two linear-mixed-model GWA tests, first to assess whether GWA could identify the large effect TSR locus (EPSPS), and the next to discover NTSR-related alleles genome-wide. The first model had only raw resistance ratings as the response variable, with no covariates. The second model included two genetic covariates: a TSR mutation of proline to serine at codon number 106 in *EPSPS* (P106S), and the magnitude of the copy number of *EPSPS* (as characterized in Kreiner et al., 2019). We included these genetic covariates to ensure that variance in resistance explained by TSR mechanisms was not attributed to other alleles across the genome and that differences in the genetic architecture of resistance across populations does not lead to spurious associations with population structure (as in Atwell et al., 2010; Sasaki, Köcher, Filiault, & Nordborg, n.d.). We did so in GEMMA (Zhou & Stephens, 2012), using the lmm-4 option and after estimation of a kinship/relatedness matrix (-gk), with covariates specified (-c option) when relevant. We employed both a Bonferroni correction and an FDR approach to significance testing in R through the function p.adjust (method=”bonferroni” or “fdr”), where we used a false discovery rate of *α* = 0.05. To calculate the significance thresholds, we took the raw, −log_10_(p-value) equivalent to a q-value (FDR corrected p-value) < 0.05. While we show both, we used the FDR cutoff for significance testing as Bonferroni correction can often be overly conservative in a GWA setting due to the assumption of independence between tests (Johnson et al., 2010), especially true in our case as we chose to not implement LD thinning.

We evaluated enrichment of and the roles of our significant SNPs (after controlling for relatedness and multiple test correction, FDR at the level of *α* = 0.05) in GO molecular functions, biological processes, cellular components, and protein classes. To do so, we identified *Arabidopsis thaliana* orthologues in the annotated female *Amaranthus tuberculatus* reference genome using OrthoFinder (Emms & Kelly, 2015) (as in Kreiner et al., 2019). Genes with significant SNPs where no *A. thaliana* orthologs were identified through orthofinder were manually curated from the reference genome annotation file (which was curated with annotations from several species to identify known genes in Kreiner et al., 2019). Finally, the set of manually curated and OrthoFinder *A. thaliana* orthologues were used to evaluate molecular function and test for enrichment of particular gene classes in Panther GO-Slim. Enrichment of significant SNPs for all Panther categories was analyzed using a Fisher’s exact test and FDR correction.

### GWA permutations

To generate a null distribution of expected significant hits, and as a point of comparison for downstream population genetic analyses for the two covariate NTSR model, we randomized phenotype with respect to multilocus genotypes. We randomized the phenotype of interest and covariates together with respect to SNP multilocus genotypes, such that the associations between the resistance and covariates were maintained—this provides an expectation for ascertainment bias of significant SNPs in regions of low recombination as well as avoids circularity between GWA ascertainment and downstream population genomic analyses. Furthermore, to incorporate the effects of population structure that may have gone uncontrolled by the kinship matrix we performed this joint randomization in two increasingly stringent ways; within geographic regions (Ontario Natural Populations, Essex, Walpole, Illinois, Missouri; where the main axis of population structure has been shown to differ; Kreiner et al., 2019), and more rigorously, within individual populations. Therefore, our empirical null distribution represents the expected results from a GWA with our given phenotypic distribution among populations, stratification between populations, and after controlling for individual relatedness.

For downstream analyses of evidence of selection, while we present distributions of significant SNPs for both permutations, our tests of significance are performed against the more rigorous within-population permutations. To do so, for each permutation type, we generated 250 sets of randomized phenotype/covariate to genotype data, running a GWA on each. To generate null distributions of the expected number of significant SNPs, we used the significance threshold (q value < 0.05) observed in the two covariate GWA model. Based on the number of significant SNPs extracted from each randomized iteration, we then calculated the average number of significant SNPs across all 250 permutations to get an ‘empirical’ FDR. For illustrative purposes we plotted the distribution of various statistics for all significant SNPs across permuted GWAs for both the within-region and within-population permutations (i.e., in **Fig 3**), and so it is important to note that these illustrated distributions reflect very different sample sizes (~500 for the observed set and ~60,000 for the permuted set). For rigorous significance testing, we relied on the more stringent within-population permutations, resampling the total pool of significant SNPs across all permuted GWAs to the same number of observations we observed in our true GWAS, 1000 times. For each resample, we took the median (or mean) of a given statistic, and then the 5 and 95% quantile to attain the 95% CI of the empirical null distribution. To test whether there was enrichment compared to the null expectation, we then compared these CIs to the median (or mean) of these same statistics for the observed GWA. We also investigated the extent to which significant SNPs from each within-population permutation was enriched for biological classes after bonferroni correction, as described above for our observed GWA.

### Summary statistics

To infer patterns of selection on putative NTSR loci and characteristics of the genetic architecture of resistance, we estimated a variety of population genomic statistics on significant hits. Allele frequency and effect sizes were analyzed directly from the output of the GEMMA GWA. Since linked selection should reduce diversity at 4-fold degenerate coding sites (putatively neutral) relative to 0-fold degenerate coding sites (putatively selective) (Smith & Haigh, 1974), we estimated diversity at these sites independently in 10 kb genomic windows around focal alleles. We used scripts from https://github.com/simonhmartin/genomics_general to centre windows on putative NTSR and permuted significant (false positive) SNPs, including both variant and invariant 0-fold or 4-fold sites. For both visualization and statistical resampling, we required windows to have data for at least 0.01% of the window (10/10000 invariant or variant sites). Tajima’s D (a summary of the extent of neutral evolution) was calculated at 0fold and 4fold sites and was estimated on a per-SNP basis, and median genomic windows were estimated with custom dplyr functions.

We also calculated three related haplotype-based statistics—extended haplotype homozygosity (EHH; Sabeti et al., 2002), integrated haplotype homozygosity (iHH), and the integrated haplotype score—that can be more powerful than single SNP or windowed approaches for identifying positive selection as they draw power from signals of selection at linked sites. All of such statistics are calculated with respect to the alternate (1) and reference (0) allele haplotypes in selscan (Szpiech & Hernandez, 2014), and so to make these statistics specific to resistant versus susceptible comparison, we swapped allele assignment (1 versus 0) only for SNPs that had a negative effect on resistance, such that all 1 alleles correspond to alleles with positive effects on resistance. These calculations require phased haplotypes and a genetic map, and therefore we called phased haplotypes in SHAPEIT (Delaneau, Zagury, & Marchini, 2013), and estimated the population-based LD map using LDhat (McVean & Auton, 2007) (as in Kreiner et al., 2019). iHS calculations build on the iHH (integrating the space under the EHH curve) by taking the difference in iHH among haplotypes and standardizing for allele frequency in bins across the genome, where haplotypes are differentially defined based on where EHH becomes < 0.05. To encompass all significant SNPs, we lowered the minor allele threshold to 0.01, as opposed to the default 0.05. Finally, we ran the program norm, implemented in selscan, to obtain the standardized iHS, which takes into account the genome-wide expected iHS value given a certain allele frequency.

All custom scripts are available at https://github.com/jkreinz/NTSR-GWAS-MolEcol2020.

## Results

### SNPs and Traits

Our final SNP set from 155 *A. tuberculatus* individuals included 10,279,913 SNPs across 16 chromosome-resolved scaffolds. Of these, 8,496,628 SNPs were used by GEMMA, after sites with > 5% missing data and an allele frequency <1% were removed. The final dataset encompassed individuals with a range of resistance levels, *EPSPS* copy number, and the EPSPS Pro106Ser substitution (Fig 1). Out of all 155 individuals, 81 were classified as glyphosate resistant (a rating of ≥2 on a scale of 1 to 5, i.e., less than ~80% plant injury) and 74 as susceptible. Note that the 1-to-5 scale was used as input for the GWA tests. Only ten out of 155 individuals had a TSR mutation in the *EPSPS* coding sequence, while the *EPSPS* amplification was much more prevalent: 79 individuals had a copy number ≤1.5, while the copy number in 76 individuals exceeded 1.5 (medians of 1.13 and 5.95 copies, respectively), with one individual ranging up to 30 copies of *EPSPS*. Seventeen individuals have no evidence for either TSR mechanism but were rated as resistant (Fig 1A;“Res., noTSR”), resistance being most likely driven solely by NTSR mechanisms. Moreover, a regression of resistance on both TSR mechanisms (EPSPS Pro106Ser and *EPSPS* copy number) explained only 33% of the variance in resistance ratings (r^2^ = 0.33, *F*_2,152_ = 38.69, *p =* 2.62e-14), implying that NTSR mechanisms (Fig 1B “NTSR”) may be widespread even across individuals with at least one TSR mechanism, which were found in 79% (64/81) of our resistant samples. Thus, genome-wide loci may be contributing both to resistance in the absence of TSR, and to the degree of resistance conferred by individuals with a TSR mechanism. In fact, because some samples with TSR were not phenotyped as resistant, coexistence of NTSR mechanisms may be required for robust phenotypic resistance.

**Figure 1.**
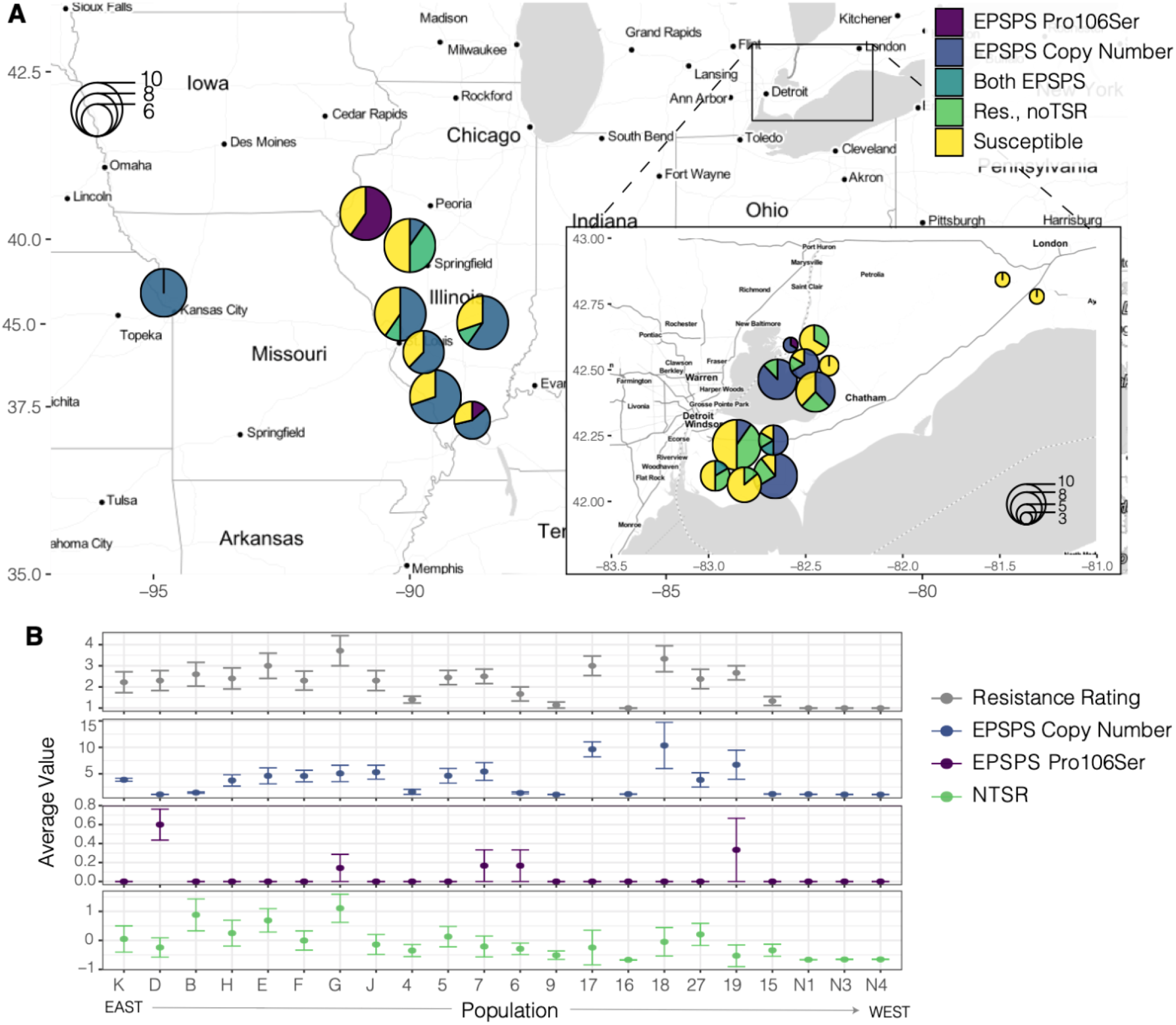
Distribution of glyphosate resistance types across populations. A) Map of populations included in the GWA, showing the proportion of susceptible and resistant individuals of different underlying genetic mechanisms based on presence/absence. “Res., noTSR” refers to individuals rated as resistant at least 2 on a 1-to-5 scale but that lack evidence for one of the two known TSR mechanisms. B) Population-means of resistant types. The NTSR rating represents the residuals from a regression of resistance on both TSR mechanisms (*EPSPS* copy number and *EPSPS* Pro106Ser), where the lowest values on the y-axis are completely susceptible. Error bars represent standard deviation of the population mean estimate.

### Gemma Linear Mixed Models & PVE

We performed two GWA mixed effect models, both of which included a random effect kinship matrix but differed in the presence of any fixed effect, genetic covariates (*EPSPS* Pro106Ser and *EPSPS* Copy Number, Table 1). The no-covariate model was run to determine whether a GWA for our resistance ratings could accurately identify the large-effect target-site resistance gene, *EPSPS*, while the two covariate model (NTSR model) was run to identify additional candidate regions across the genome contributing to polygenic resistance, controlling for any linkage effects with TSR and/or differences in the genetic architecture of resistance among populations that might lead to spurious associations between TSR and population structure. The no-covariate model had a lambda of 0.979 and the qq-plot showed a large tail of an excess observed-to-expected p-values. While the elevated tail in the qq-plot was less pronounced for the NTSR two-covariate GWA model, the lambda estimate was the same, and this model was associated with a higher maximum likelihood estimate (Table 1, Sup Fig 1).

**Table 1.** GWA results from two linear mixed models varying in their fixed effect covariates, based on the same genome-wide SNP set & kinship matrix as a random effect.

| Covariates | $-\log_{10}(p)$<br>at $q = 0.05$ | # Sig.<br>SNPs<br>@<br>FDR | PVE<sup>†</sup><br>(SE) | MLE | Lambda<br>(95% CI) | Global<br>Permuted<br>FDR<br>estimate | Within-Region<br>Permuted<br>FDR estimate | Within-Pop<br>Permuted<br>FDR<br>estimate |
| --- | --- | --- | --- | --- | --- | --- | --- | --- |
| None | 6.4 | 91 | 0.388<br>(0.230) | -268 | 0.979<br>(0.972,<br>0.986) | 0.011<br>(1/91) | 0.04<br>(4/91) | 0.06<br>(5/91) |
| EPSPS<br>TSR SNP +<br>Copy Num. | 5.6 | 573 | 0.374<br>(0.219) | -234 | 0.979<br>(0.971,<br>0.986) | 0.096<br>(55/573) | 0.29<br>(165/573) | 0.40<br>(229/573) |

<sup>†</sup> when covariates are included, PVE represents percent of *residual* variance explained

The no-covariate model had 91 significant SNPs after correction, compared to 574 SNPs in the two covariate, NTSR model. The global-, regional-, and population-level permutations provided estimates of the ‘empirical’ FDR based on the average number of SNPs that are significant at this threshold across 250 phenotype-genotype randomized GWAs. Despite using a stringent FDR cutoff of *α* = 0.05 for significance testing in both the covariate-free and two-covariate model, we found with these permutations relatively large numbers of significant SNPs (55, 165, and 229 out of the 573, respectively) (Table 1). The number of significant SNPs in permutations suggested a residual effect of population structure unaccounted for by the initial significance cutoff and controlling for the kinship matrix—or put another way, that differences in the genetic architecture of resistance among populations covary with fine-scale population structure. While these empirical estimates of the FDR highlight that a number of these SNPs we characterize here may result from either of these processes and individually should be treated with caution, we examine evidence for selection and relevant biological functions against the most stringent within-population permuted set, and provide a conservative estimate for the amount of variance in resistance that can be explained by these putatively NTSR-related alleles.

The relative importance of monogenic versus polygenic mechanisms of resistance can be explored by comparing the percent of variance explained (PVE) by genome-wide SNPs in the two covariate NTSR model (polygenic effects on resistance) to the r^2^ of glyphosate resistance on TSR and EPSPS copy number (monogenic effects). Controlling for both TSR genetic mechanisms leads to a marginal PVE estimate of 0.374, implying that genome-wide SNPs can explain 37% after accounting for resistance associated with TSR at EPSPS. The initial multiple regression model of resistance on monogenic mechanisms (TSR SNP + EPSPS copy number) results in an r^2^ = 0.33 (*F*_2,152_ = 38.69, *p =* 2.62e-14). Thus, a rough estimation of the residual variance unexplained by TSR mechanisms (1-0.33 = 0.67) suggests that the NTSR loci we identify here can explain about a quarter (0.37 x 0.67) of the variance in resistance—implying that even in a sample of resistant individuals that nearly all segregate for glyphosate TSR, NTSR mechanisms may be of near equal importance.

We also implemented a multiple regression of resistance ratings on both TSR genetic covariates, including the kinship matrix as a random effect covariate to estimate the SNP heritability of resistance, finding 83% of variance in resistance can be explained by additive genetic effects. This provides further support for the argument that a significant component of the variation in resistance that is unexplained by TSR is due to loci distributed throughout the genome.

### Distribution of genome-wide associations, candidate genes, and GO enrichment

The qualitative pattern of GWAs with glyphosate resistance differed greatly between the no-covariate and NTSR, two covariate models (Fig. 2a,b). In the no-covariate model, a butte of significant SNPs within the confines of the EPSPS-related amplification is apparent on scaffold 5 (Fig. 2b) with glyphosate resistance associated loci found directly within EPSPS. In contrast, the NTSR two-covariate model shows no signal of association at EPSPS, as is expected given our EPSPS-based covariates. The NTSR model shows significant hits distributed across all 16 chromosomes, with particularly high concentration of significant hits on scaffold 1, 15, and 2 (42%, 18%, 10% of genome-wide significant hits, respectively) **(Fig 2b,c)**. Importantly, a number of genome-wide significant hits found in the no-covariate model become non-significant in the NTSR model. We interpret this to reflect the fact that population-level differences in the genetic basis of resistance (**Fig 1**) can lead to spurious genome-wide associations due to covarying population structure. A potential confounding of population structure and the genetic basis of resistance highlights the importance of the two-covariate model, as well as the regional and within-population permutations, for identifying variants associated with NTSR.

**Figure. 2.**
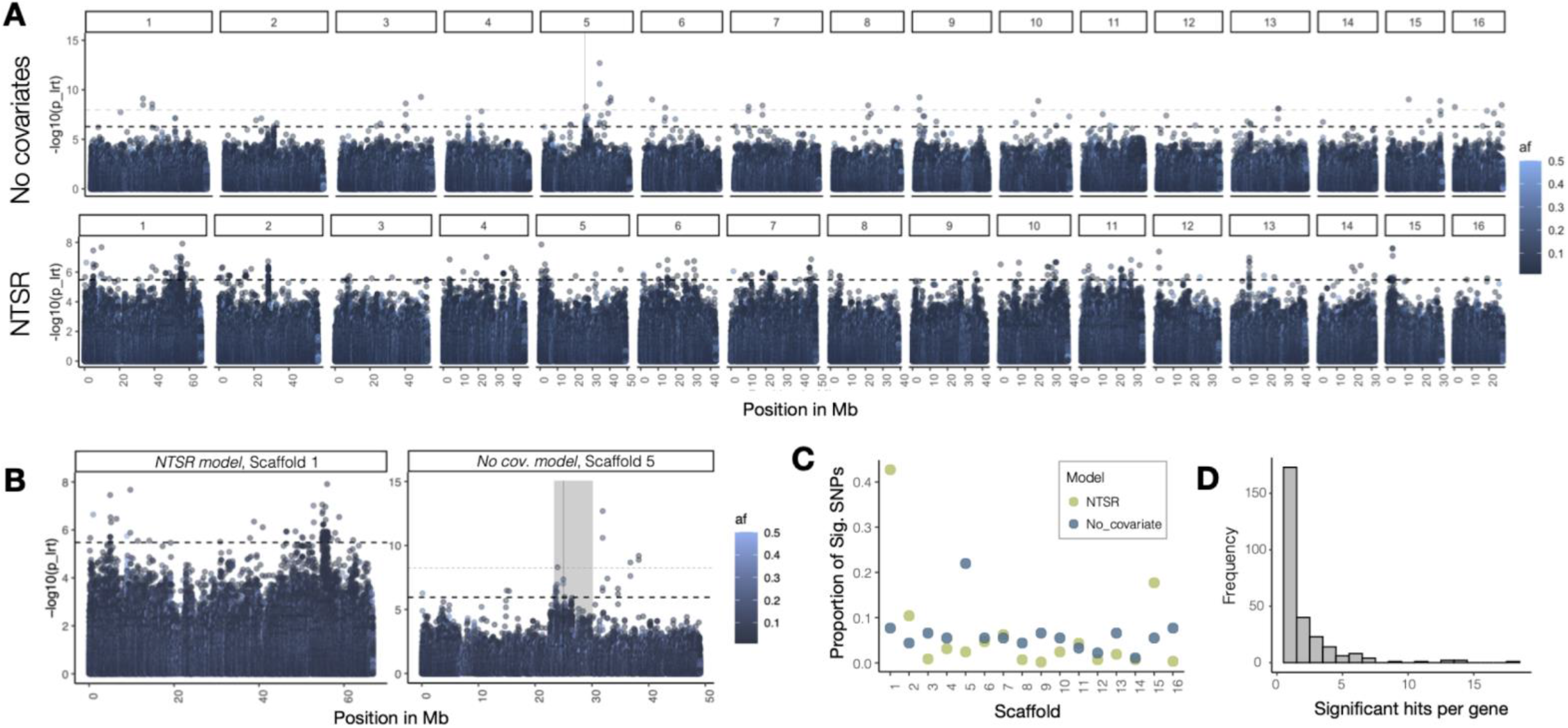
GWA of glyphosate resistance. A) Manhattan plot of significance of association with glyphosate resistance across all 16 scaffolds for two linear mixed models - the upper without fixed effects and the lower with TSR as a covariate. Dashed black horizontal bar indicates the 0.05 FDR threshold, while the dashed grey bar above indicates the Bonferroni correction threshold. Thin vertical line on scaffold 5 depicts the EPSPS gene. X axis is scaled by the length of each scaffold. B) Manhattan plot for scaffold 1 (containing 42% of significant hits) [Left] as compared to the Manhattan plot for scaffold 5 in the no covariate model, which contains the target gene for glyphosate, EPSPS [Right]. The vertical grey rectangle indicates a previously identified ~6.5Mb EPSPS amplification present across individuals in this dataset. C) The proportion of significant SNPs present on each scaffold for each linear mixed model. D) The number of significant hits per gene in the NTSR model.

The set of 573 significant SNPs with −log10(p) values corresponding to q < 0.05 (Table 1) in the NTSR model corresponded to 274 unique *A. tuberculatus* genes. Six genes have more than 10 significant SNPs mapping within them, four of which are found on Scaffold 1 in a peak between 54.9Mb and 56.3Mb (Fig 2b,c). These four genes are a DNA-directed DNA polymerase (AT5G44750), DNA-directed RNA polymerase family protein (AT2G15430), B-block binding subunit of TFIIIC (AT1G59077), and Auxin-responsive protein SAUR21 (AT5G18030) (Fig 2b). On Scaffold 2, a Vacuolar protein sorting-associated protein 26A (AT5G53530) also has >10 SNPs mapping.

For 150/274 of these *A. tuberculatus* genes, we could find mappable orthologs in *Arabidopsis thaliana*, and thus could be used as input for enrichment analyses and candidate gene exploration. Fisher’s exact test showed this set of genes to be significantly enriched after FDR correction for 13 distinct classes of GO biological processes in 6 hierarchical categories **(Fig 3)**, the two most significant independent terms being response to chemicals and biosynthetic processes.

**Figure 3.**
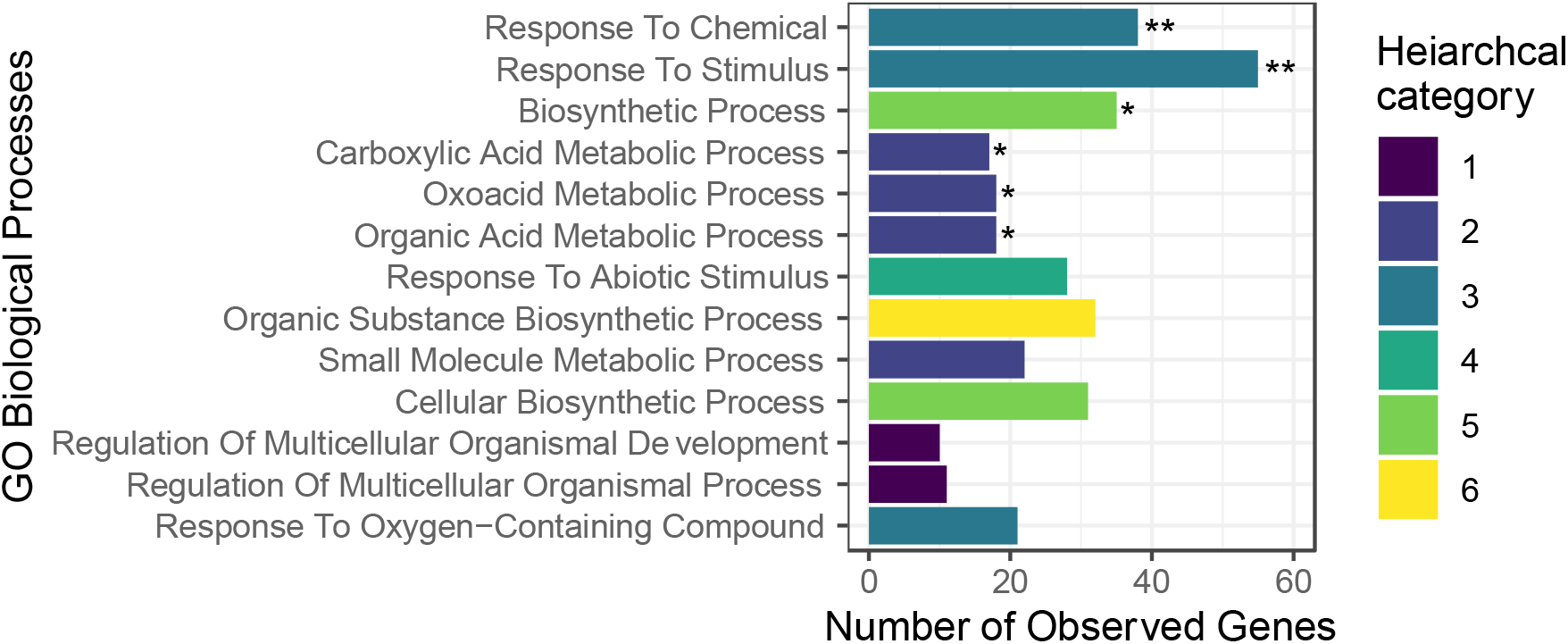
573 SNPs significantly associated with NTSR are enriched for 13 GO biological processes after FDR correction. Classes are sorted by increasing raw p-value, and coloured by hierarchical categories. Categories significant after Bonferroni correction are indicated by asterisks (*** = p-value < 0.0001, ** < 0.01, * < 0.05).

Many of the biological processes of genes in our set of significant SNPs were notable given previous work on particular gene families and the three known molecular phases of NTSR (Gaines et al., 2020) (1 - herbicide detoxification via oxidation, hydrolysis, or reduction, 2 - conjugation, and 3 - compartmentalization/transport) and metabolic pathways known to be modulated by the Shikimate pathway targeted by glyphosate. The 573 alleles we found significantly associated with NTSR function in relevant biological processes such as response to chemicals, stimulus, and response to oxygen-containing compounds, and numerous metabolism and biosynthetic and metabolic related processes (Fig 3). The highest proportion of NTSR-related genes function broadly in response to stimulus (55/150) or chemicals (38/150), and biosynthetic processes (35/150), all of which remain significant even after stringent bonferroni p-value correction, along with 3 other carboxylic acid metabolic process-related classes (Fig 3). In comparison, across the 250 within-population GWA permutations, 225 have zero GO biological classes withstanding bonferroni correction, with the remaining iterations having only 1.17 terms passing bonferroni correction on average. Furthermore, none of these GWA iterations are significant for any of the GO biological processes we observe to pass bonferroni testing.

Key genes involved in response to chemicals and organic acid metabolic processes code for many protein classes, including aldolases (FBA6, FBA2 - regulate glycolytic processes in response to abiotic stress (Lu et al., 2012)), a dehydrogenase (CICDH - NADPH dependent dehydrogenase that plays a key role in redox homeostasis (Mhamdi et al., 2010)), decarboxylases (ADC1), kinases (BSK2, BAM1, PI4KG4, ADK1, LIP1), reductases (GLO5, GRXC1 – involved in redox regulation (Riondet et al., 2012)), methyltransferases (JMT - involved in plant defense, induced by wounding and methyl-jasmonate treatment (Seo et al., 2001)), WRKY transcription factors (WRKY27, WRKY1; known to play a key role in biotic and abiotic stress response (Pandey & Somssich, 2009; Phukan, Jeena, & Shukla, 2016)), auxin responsive genes (SAUR5; SAUR50, DTX51; EXO70A1, ARF1), ethylene receptors (ETR1,ERF061), and an ABC transporter (ABCG39, part of the pleiotropic drug resistance (PDR) transporters, ABCG subfamily). A full list of Amaranthus and Arabidopsis genes corresponding to SNPs significantly correlated with NTSR can be found in supplement file 1.

While many of our NTSR-associated genes function in metabolic and chemical response pathways, we also see enrichment for categories less obviously related to detoxification— notably, regulation of multicellular organismal processes and development. Examples of the 11 genes enriched for these developmental processes include: *GIGANTEA*, encoding a nuclear protein with pleiotropic roles in flowering time, stress response, circadian clock regulation, with prior evidence for providing herbicide tolerance by increasing oxidative stress resistance (Kurepa, Smalle, Va, Montagu, & Inzé, 1998; Mishra & Panigrahi, 2015); *EMBRYONIC FLOWER 2* (*EMF2*), which in *A. thaliana* prevents premature flowering (Yoshida et al., 2001), *PETAL LOSS* (*PTL*), which regulates perianth architecture (Brewer et al., 2004); *BBX18,* which suppresses photomorphogenesis by blue-light mediated hypocotyl reduction (Vaishak et al., 2019; Wang et al., 2011), and *UNFERTILIZED EMBRYO SAC10* (*UNE10*) and *ARM REPEAT PROTEIN INTERACTING WITH ABF2* (*ARIA)*, two negative regulators of seed germination (Tepperman et al., 2004; Kim et al., 2004).

### Genomic context of putative NTSR alleles

With a better understanding of the types of genes putatively involved in NTSR, we posited that a closer look at the genomic context of the loci in our set of candidate genes would shed light on the action of selection, genetic architecture, and heterogeneity of NTSR among populations. To do so, we compared various characteristics of our observed NTSR-related SNPs to the within-population permuted expectation, such that particularly extreme values indicate evidence above and beyond the effects of genetic and phenotypic population structure or recombination heterogeneity across the genome.

Our reference genome was from a glyphosate-susceptible individual, and we polarized our SNPs against this reference. The effect sizes of non-reference alleles were highly asymmetric in direction; of the 573 non-reference variants, only 8 SNPs had negative effects on glyphosate resistance compared to 565 with positive effects. Our significant SNPs broadly fell into two categories, rare alleles of modest to large effect and common alleles with small effects (Fig. 3A). The negative relationship between effect size and frequency may be due to biological or methodological properties, or a combination of the two: resistance alleles with larger effects may have more deleterious pleiotropic effects (as in Kryukov, Pennacchio, & Sunyaev, 2007; Marouli et al., 2017; Tennessen et al., 2012, Josephs, Lee, Stinchcombe, & Wright, 2015), a lack of power makes it difficult to detect rare, small-effect alleles, and both the Winner’s Curse and Beavis effect lead to an overestimation of detected effect sizes (Liu & Leal, 2012; Xu, 2003). The tradeoff between effect size and allele frequency was clearly present in the genetic architecture of NTSR, where our set of significant SNPs were enriched for large absolute effect sizes compared to the empirical null distribution (median beta = 2.455; [95% CIs of median empirical null distribution, based on resampling: 2.084, 2.186]), and for rare alleles (median AF = 0.016, [95% CIs of median empirical null: 0.019, 0.023]) (Fig. 4A,B).

**Figure 4.**
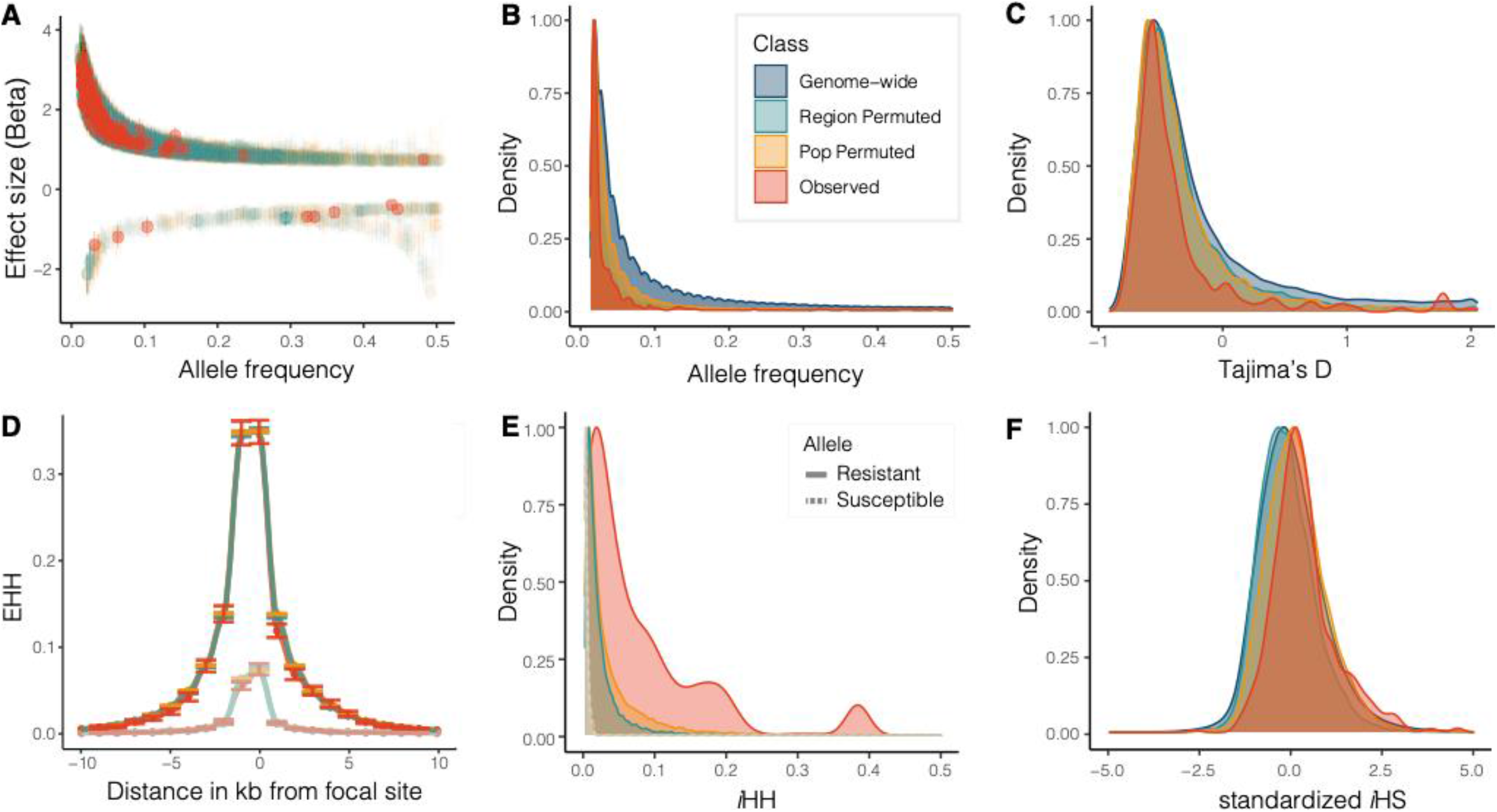
Genomic summaries of SNPs significantly associated with NTSR compared to the empirical null distribution and genome-wide background. A) Effect sizes relative to allele frequencies, B) the distribution of allele frequencies, C) the distribution of Tajima’s D, D) expected haplotype homozygosity (EHH) around focal resistance-related and susceptibility-related alleles, E) the integrated haplotype homozygosity of resistant-related versus susceptibility-related haplotypes, F) standardized integrated haplotype score (iHS), treating resistance alleles as the derived allele (note a positive value indicates an excess of derived relative to ancestral haplotype size, as in Szpiech & Hernandez, 2014). The legend in the B applies to all plots, while legends for D and E additionally distinguish allele type.

We tested for signals of directional selection on NTSR-related alleles by examining the length and homozygosity of resistant versus susceptible haplotypes. We first investigated signals of extended haplotype homozygosity (EHH), a haplotype-based test that has been used to assay signals of positive selection around focal SNPs based on decay of homozygosity. When we truncated the EHH distribution to 10 kb around either side of observed resistance-associated alleles, we saw negligible differences in average EHH value across this region in observed resistance-, compared to permuted resistance-associated haplotypes (median R allele EHH across 20kb = 0.000473, 95% CIs of mean empirical null: 0.000, 0.0196) (Fig 3D). Taking the integral of the EHH by distance curve referred to as integrated Haplotype Homozygosity (iHH; Voight, Kudaravalli, Wen, & Pritchard, 2006)—but without truncating haplotype length—revealed that observed resistant haplotypes were longer than expected under the empirical permuted null distribution (median iHH = 0.042531, [95% CIs of median pop-permuted empirical null: 0.01552, 0.01929]) (Fig 3E). Finally, the standardized integrated Haplotype Score (iHS) (which compares differences in iHH between reference and ancestral alleles, standardized for allele frequency; Voight et al., 2006) identified enrichment for unusually large homozygous tracts of resistant relative to susceptible haplotypes, given their respective allele frequencies (median standardized iHS = 0.21286, [95% CIs of median pop-permuted empirical null: 0.00785, 0.13600]) (Fig 3F).

These haplotype-based inferences suggest a role for recent directional selection, and so as a final check we also estimated diversity at Tajima’s D at 0-fold and 4-fold degenerate coding sites in 10 kb windows surrounding focal SNPs, where signatures of linked selection should show particularly strong signals of selection at putatively neutral 4-fold sites. We see no evidence for reduced 4-fold diversity or 0-fold diversity around NTSR-related SNPs compared to the permuted expectation (median 4fold *π* = 0.0439, [95% CIs: 0.0415, 0.0447]; median 0fold *π* = 0.0152759 [95% CIs: 0.01528749, 0.01728545]). However, whereas the observed SNP set had sufficient data (at least 10 variant or invariant degenerate sites of a given type) in 419/573 4-fold and 443/573 0-fold windows (therefore representing ~75% of relevant windows) the population permuted set had only 784/78036 and 814/78036 (~1%), limiting the inferences that can be drawn from this comparison. We see more negative Tajima’s D than expected in windows of 4-fold but not 0-fold sites (median 4fold Tajima’s D = −0.533319 [95% CIs: −0.482590, − 0.429297]; median 0fold Tajima’s D =-0.5902975, [95% CIs: −0.606059, −0.571910]), (Sup Figure 2). Lastly, we find our observed GWA SNP set is enriched for polymorphic 0-fold degenerate coding sites relative to the empirical null distribution (proportion of 0-fold sites = 0.0925%, [95% CI of the proportion of 0-fold sites per GWAS itt = 0.0137, 0.0728]).

## Discussion

Herbicides, when appropriately applied in agricultural settings, are often lethal, and for this reason they elicit very strong and rapid evolutionary responses in weed populations. The most commonly recognized genetic responses to these strong selection pressures are the appearance of mutations at the loci whose products are directly targeted by the herbicide. How often selection by herbicides leads to easily detectable changes at other loci in the genome has remained unclear. In this study, we investigated the genetic architecture of resistance to glyphosate herbicides in a problematic agricultural weed, *Amaranthus tuberculatus.* As proof of concept, GWA correctly identified *EPSPS*, the target gene of glyphosate herbicides, and many significant SNPs neighbouring this region as significantly associated with resistance. This suggests considerable LD around EPSPS, consistent with previous findings of a strong selective sweep associated with a ~6 Mb region that is amplified (Kreiner et al., 2019). Consistent with polygenic NTSR being widespread across populations and the genome, however, we find that after controlling for variation in resistance explained by TSR loci, SNPs in more than 250 distinct genes across all 16 chromosomes are also associated with glyphosate resistance.

Compared to both the genome-wide background and null-expectation, we find that NTSR-related alleles are enriched for particularly low frequencies and large effects. Together, these results fit a scenario of a genetic architecture of NTSR shaped by recent selection but also strong pleiotropic costs, with the involvement of specific NTSR-related alleles being highly heterogeneous across populations and genetic backgrounds. Below, we discuss the genetic architecture of glyphosate resistance, the role of selection in shaping it, and candidate NTSR genes.

### Model comparison, importance of NTSR versus TSR mechanisms, and population structure

With previous knowledge of two genetic causes underlying glyphosate resistance—a non-synonymous mutation within *EPSPS* that prevents glyphosate from inhibiting the encoded enzyme, and amplification of *EPSPS* that overcomes the inhibitory effect of glyphosate through overproduction of the EPSPS protein—we sought to control for variance in resistance explained by TSR to identify genome-wide SNPs contribution to polygenic, NTSR (Table 1). Reassuringly, the covariate free model showed enrichment for significant SNPs on the scaffold harbouring *EPSPS* (with 32 SNPs within the *EPSPS*-amplification significantly related to glyphosate resistance and ~20% of significant GWA SNPs located on this scaffold), while controlling for TSR mechanisms led to the *EPSPS*-containing scaffold having a similar number of significant hits as an average chromosome (Fig 2).

Our data encompassed a set of 155 individuals where 52% are phenotypically resistant, 79% of which harboured at least one large effect TSR mechanism. That 25% of phenotypic resistance can be explained by polygenic NTSR in this sample where large-effect TSR mutations are so widespread, explaining a comparable 33% of variance in resistance, speaks to the underappreciated importance of polygenic NTSR—which may not only confer resistance in the absence of TSR, but may also act as modifiers of resistance in the presence of TSR. While this pronounced importance of polygenic resistance we describe here may even be a conservative estimate if highly resistant individuals that lack TSR mechanisms are more widespread in the wild than represented in our sampling, underpowered GWAs are known to overestimate the PVE/”chip heritability” (King & Long, 2017), a source of bias which we cannot rule out here. Furthermore, given our estimate of the heritability of resistance being 0.83, it is evident that the variance explained by TSR (0.33) and NTSR mechanisms (0.25) does not fully resolve the genetic basis of resistance.

Among populations, the importance of NTSR and TSR mechanisms are bound to vary as the involvement of these genetic mechanisms differ across populations, geographic ranges, and species (see Gaines et al., 2020 for a thorough review). This is evident in our results (Fig 1), where the mechanisms and presence of TSR vary considerably even across nearby populations. When focal phenotypes, or its genetic architecture, differ among populations in a manner related to genome-wide and fine-scale relatedness among populations, alleles that come out as significantly related to the trait of interest may be confounded with shared evolutionary history. A relationship between population genetic structure, genomic background, and the genetic mechanism(s) of a trait are a key limitation inherent to most GWAs that sample across a diverse set of populations, in that this signal may go uncontrolled by a relatedness matrix (discussed in Atwell et al., 2010). With this limitation in mind, we examined the extent that the genetic architecture of resistance differed among populations in a manner that might be confounded with population structure.

To do so, we took three increasingly rigorous approaches. We (i) permuted the GWA 1000x, randomizing phenotypes and covariates with respect to genotypes globally among all samples, addressing the problem of circular downstream population genomic analyses by having an empirical null that should be equally affected by such circularities. However, to do so in a way that can shed light on the extent to which the genomic architecture of non-target site resistance may be confounded with population structure, we (ii) performed these permutations within geographic regions that were previously found to be the main axes of differences in population structure in this dataset (Kreiner et al., 2019), and (iii) performed an even more rigorous permutation within populations. While the global permutations find on average many fewer significant SNPs than our observed set (55/573), our regional permutations and population permutations increasingly find a large proportion of SNPs significantly related to NTSR even under these randomized set of phenotypes (165 and 229/573, respectively). Thus, it is evident that residual population structure (unaccounted for by our relatedness matrix) and the genetic architecture of resistance are at least partially confounded, as we obtained a large number of significant SNPs even when phenotypes were randomly assigned to genotypes.

To gain some intuition behind these within population permutation results, consider a situation in which resistance phenotype, TSR genotypes, and SNP identity at a hypothetical NTSR site in the genome are highly correlated, or even where there is only 1 or limited combinations of genotypes or genomic backgrounds in a population. In these instances, randomly assigning phenotypes to different genomic backgrounds may not change the p-values or test statistic of a GWA, as each phenotype is being randomly assigned to the same multilocus genotype, albeit from a different individual. Consequently, under many permutations, we will observe p-values or test-statistics equal to the original observation, despite random permutation, and these statistical observations may reflect loci that have experienced selection from herbicides in relatively homogeneous populations. Alternatively, polygenic NTSR-based resistance can become associated with genome-wide SNPs that tag local population structure, even if they are not causally associated with resistance.

While our candidate genes will need to be carefully probed in other sets of populations and with other experimental approaches, the set of NTSR-related alleles we describe here show particularly extreme signals of selection and herbicide resistance-related biological functions, even against the most stringent population-permuted empirical null model that accounts for uncontrolled population structure. Our results speak to widespread nature of polygenic resistance in *Amaranthus tuberculatus* agricultural populations in North America—not just supplementing low levels of protection against a background of large-effect TSR mechanisms, but providing near equivalent provision of protective effects against glyphosate. Further sampling would also provide an interesting comparison of the extent to which the genetic architecture of NTSR is population-specific, and how well components of our analysis are reproduced in a set of samples with markedly different population structure and possibly different architectures of resistance (as has been discussed in Josephs, Berg, Ross-Ibarra, & Coop, 2019).

### Effect size & Allele frequencies

The effect sizes of SNPs associated with NTSR tend to be negatively correlated with allele frequency (**Fig 4A)**. The lack of rare alleles with small effects is likely to be driven by a lack of statistical power—a key driver in the problem of missing heritability (Manolio et al., 2009). For human height, a classic example of a polygenic trait where small effect sizes should be prevalent, only 83 significantly associated alleles with a frequency < 0.05 could be identified with a sample size of 711,428 individuals, and their effect sizes were >10 times that of common variants on average (Marouli et al., 2017). The median effect size (beta) of rare alleles (<0.05) in our GWAS was 2.71, while for common alleles (=>0.05) it was only 1.10 (only 57/573 significant SNPs). These beta values can be interpreted as the effect on resistance, per changing the count of the allele by 1. In our case, a unit of resistance on a scale of 1-5 corresponds roughly to a 20% decrease in percent damage from glyphosate application. Thus, for every increase in allele count, the median effect size rare allele may lead to a near 50% decrease in percent damage.

Aside from power, the negative correlation between allele frequency and effect size and the general lack of intermediate or common, large effect alleles may be a consequence of selection. For human diseases and gene expression in *Capsella grandiflora* and in maize, this phenomenon has been explained by greater deleterious, and possibly pleiotropic, costs of large effect alleles, such that purifying selection reduces their frequency within populations (Eyre-Walker, 2010; Josephs, Lee, Stinchcombe, & Wright, 2015; Kremling et al., 2018; Kryukov et al., 2007; Marouli et al., 2017). The enrichment for large effect alleles is likely to reflect particularly strong selection in shaping the genetic architecture for herbicide resistance (Eyre-Walker, 2010)). With herbicide resistance, selection for resistance approaches near lethality (s ~ 1), and thus the trade-off between these large effect alleles being advantageous on the short-term but associated with potentially long-term pleiotropic costs may contribute to keeping such alleles globally rare.

Similarly, the range in allele frequency and enrichment of rare alleles also implies that while a small subset of resistance-related alleles are shared among populations, most appear to be population specific. The mean frequency of NTSR alleles is 0.03 in our samples, implying such alleles are found in only 4-5 individuals. Rare population-specific alleles are likely to be the work of compensatory mutations arising on already resistant backgrounds (as is the case for antibiotic resistance; Craig MacLean, Hall, Perron, & Buckling, 2010), that interact epistatically to strengthen resistance to herbicides or mediate their costs. The stacking of relatively small-effect alleles as an important genetic mechanism of NTSR has been hypothesized many times (Délye, 2013; Gressel, 2009; Kreiner et al., 2018) and rapid evolution of resistance by diverse genetic mechanisms in just a few generations of selection have provided support for such a polygenic basis (Busi & Powles, 2009; P. Neve & Powles, 2005). Our work, finally provides some of the first evidence of the frequencies and effect sizes of NTSR alleles across populations, and that when combined, such alleles may provide higher resistance than TSR mechanisms.

### Candidate genes

We found 573 NTSR-associated SNPs, corresponding to 274 unique *Amaranthus tuberculatus* genes. Of these genes, we could annotate 150 using orthology-based approaches. We found our gene set had diverse roles in biological processes and molecular functions, with 13 GO terms significant after FDR correction. Enriched biological processes were highly relevant not to just the three-phase schema of non-target site resistance (Gaines et al., 2020)—detoxification, conjugation, and transport—but to downstream pathways known to be modulated by the shikimate pathway targeted by glyphosate. Likely our most convincing evidence that the genetic architecture we describe here is related to glyphosate usage comes from the fact that our most significantly enriched GO category is response to chemicals. The genes involved in response to chemicals are diverse in their function playing roles in other enriched classes such as biosynthetic processes, response to abiotic stimulus, response to oxygen-containing compounds, and several types of carboxylic acid metabolism. Importantly, these enrichment results are much more extreme than is seen in any of within-population GWA permutations.

In particular, a recent analysis of the transcriptomic response to glyphosate in genetically modified soybean with one or multiple stacked TSR mechanisms (Zanatta, Benevenuto, Nodari, & Agapito-Tenfen, 2020) provides an interesting case study, as the genetic architecture of NTSR we identify is predominantly on the background of two large-effect TSR mechanisms, akin to their genetically modified stacked-TSR backgrounds. The KEGG pathways in soybean with the largest number of genes involved in response to glyphosate were metabolic pathways and biosynthesis of secondary metabolites (Zanatta et al., 2020). In comparison, of our 6 GO categories that were significant even after conservative Bonferroni correction, 4 were related to metabolism or biosynthesis. In addition to these broad metabolic pathways, our NTSR genes function in carbon-, and oxygen-related metabolic processes. Increased carbon-flux and thus carbon-related metabolism may compensate for reduced levels of aromatic amino acid production in response to EPSPS inhibition by glyphosate (Maeda & Dudareva, 2012).

The synthesis of several natural products, including pigments and hormones, directly require intermediates from the shikimate pathway (Maeda & Dudareva, 2012); similar to Zanatta et al., we find that genes functioning in response to chemicals are involved in diverse regulation of auxin, ethylene, abscisic acid, jasmonic acid, and gibberellin. Since glyphosate accumulation occurs in active metabolic tissues (Cakmak, Yazici, Tutus, & Ozturk, 2009), the primary location of plant hormone production, part of the toxicity induced by glyphosate is known to be the hormonal imbalance that influences both growth and development (Gomes et al., 2014). Relatedly, our genes were enriched for function in multicellular organismal processes and development, with roles in dormancy, germination, photomorphogenesis and flowering time, several of which also being enriched in response to herbicides in the single-transgene soybean variety (Zanatta et al., 2020). Together, our results suggest a key role of pleiotropic constraints, in the form of trade-offs with growth and reproduction, mediating the frequency of alleles contributing to the polygenic bases of glyphosate resistance.

While pleiotropy may be one explanation for why we see genes with diverse functions in our GWA for glyphosate resistance, and this would not be unexpected given our understanding of the organismal-level impacts of herbicides (Comont et al., 2019; Dyer, 2018; Kuester, Fall, Chang, & Baucom, 2017), a complex history of compounded selection from various herbicides and related shifts in life history optima could also contribute to this pattern. The problem of correlated traits is endemic to all GWA style analyses (Coop, 2019; Novembre & Barton, 2018; Racimo, Berg, & Pickrell, 2018), which is not a new problem in evolutionary genetics (Lande, 1979; Lande & Arnold, 1983). GWA studies are inherently correlative, with traits (response variables) related to SNP genotypes through a statistical model. It is important to keep in mind that these types of analyses, in addition to describing the genetic basis of the response variable, can also capture correlated traits in at least two ways. First, after the origin of TSR resistance, it is likely that any trait that improves plant performance in the presence of glyphosate damage (e.g., growth rates, generalized stress responses, changes to photosynthetic performance, water relations, phenology, etc) will be favored, even if these traits are physiologically and pleiotropically unrelated to detoxification, conjugation, or transport of herbicides. Consequently, SNPs that improve these traits will be associated with phenotypic assays of resistance and observed as GWA hits. Second, any agricultural practice (pesticide use, manuring, tilling, or irrigation), soil characteristic, or climate variable that is correlated with glyphosate usage and thus resistance ratings will create a set of co-selected traits (e.g., resistance, nutrient uptake, seed germination dormancy, flowering time, drought tolerance), and these may also be detected as a GWAS hit for herbicide resistance, even if their function is for other ecologically important traits.

While we would argue that the genetic architecture of NTSR we characterize here as having convincing and direct function in response to and compensation of glyphosate toxicity, the interpretation of these resistance-related alleles as strictly driven by selection only from glyphosate herbicides is likely incorrect. These putative NTSR alleles are likely to characterize a complex history of agricultural selection in resistant weeds, and if these regimes differ among populations, we expect this complex genetic architecture to be confounded with population structure. To disentangle the complexities of how shared the genetic basis of NTSR is in response to different herbicides and with other non-resistance related traits, complementary quantitative genetic and population genomic methods could be applied to cohort of genotypes with many phenotyped traits to estimate correlated selection and corresponding genetic architectures.

### Haplotype homozygosity, and a role for recent positive selection

Our set of resistance-related alleles was enriched for large effect sizes and low allele frequencies (Fig 4A,B). Furthermore, we found enrichment for particularly long haplotypes associated with resistance given their allele frequency (Fig 4D-F), particularly negative values of Tajima’s D, and a high proportion of amino-acid changing SNPs. These particularly extreme values allowed us to interpret the history of selection on the genetic architecture of NTSR, as it is above and beyond what the permuted expectation that accounts for uncontrolled population structure in both phenotype and genotype.

Selection on genome-wide resistance-related alleles may be in the form of positive and/or balancing selection. Because net selection for and against herbicide resistance should vary depending on environment, assuming a direct or pleiotropic cost of resistance, both susceptible and resistant individuals may be alternately favored in herbicide and herbicide-free settings over many generations (spatially and/or temporally fluctuating selection). Weed science research has long been interested in the costs of herbicide resistance mutations (Lenormand, Harmand, & Gallet, 2018), and work on the topic has led to sometimes conflicting conclusions, with fitness costs dependent on the mutation, organism, and environment (summarised in Baucom, 2019). Nonetheless, studies on the costs of resistance in the absence of herbicides have focused primarily on TSR mechanisms, and pleiotropic costs of resistance as a trait should be much more ubiquitous where hundreds of diverse genes are likely to be involved (e.g. 90% of all human GWAS hits overlap multiple traits; Watanabe et al., 2019).

We find no evidence that extended haplotype homozygosity (EHH) is enriched in a distance of 10 kb around significant hits in our observed NTSR-related alleles compared to the null expectation and alternative susceptible haplotypes (Fig 4D). However, both iHH (integrated haplotype homozygosity, representing the space under the EHH curve) **(Fig 4E)** and a further standardized version of this statistic, the standardized integrated haplotype score (iHS; Fig 4F), are much larger than expected relative to the permuted null expectation. Since iHS is standardized by genome-wide empirical allele frequency distributions, this can be interpreted as unusually large homozygous haplotypes associated with resistance relative to susceptible alleles (Fig 3E) (Voight et al., 2006). Particularly long haplotypes associated with resistance are expected under partial selective sweeps, which should increase the homozygous haplotype length of selected compared to unselected alleles (Voight et al., 2006)). In contrast, balancing selection, which is expected to maintain diversity over longer timescales, should lead to particularly short tracts of homozygous haplotypes as old haplotypes have both high diversity and allelic associations broken-up over time due to more opportunity for recombination (B. Charlesworth, Nordborg, & Charlesworth, 1997; D. Charlesworth, 2006; Hudson & Kaplan, 1988). That we see an excess of both standardized iHSs and resistant iHH values implies that resistance alleles have been subject to positive selection. Furthermore, these haplotype-based analyses were consistent with enrichment for particularly low Tajima’s D in windows, and enrichment of amino-acid changing SNPs. Taken together, base-pair-, haplotypic-, and window-based summary statistics point to the genetic architecture of NTSR having been the subject of recent positive selection, constrained by pleiotropic trade-offs.

Early population genetic theoretical studies have examined the effects of rotating crop and herbicide use (a rotation scenario of one year on/one year off) on herbicide resistance, assuming selection coefficients (*s)* for resistance between 0.75 and 0.99 within generations, and a cost of resistance in the absence of herbicides as *s*=0.25. Under such scenarios, a rare monogenic dominant allele will reach a frequency of ~0.7 (*s*=0.75) or go to fixation (*s*=0.99) within 20 years (Jasieniuk, Anita L. Brûlé-Babel, & Ian N. Morrison, 1996). Our results illustrate that the genetic architecture of herbicide resistance in our system is more complex, with a polygenic basis rather than a single dominant locus, and with a range of allele frequencies and effect sizes. This complex basis means that small changes in the frequency of many alleles can appreciably alter phenotypic values, with weaker selection on each allele individually required. Compounded with a role of pleiotropic constraints, diversity in polygenic resistance-related alleles may persist in populations over much longer timescales then initially hypothesized.

## Conclusion

In conclusion, our study identifies over 500 SNPs in more than 250 genes across all 16 chromosomes associated with non-target site resistance in *Amaranthus tuberculatus*. These alleles are associated with large effect sizes but low frequencies, indicating a heterogeneous architecture of NTSR across populations. Consistent with the literature, we find NTSR genes function in pathways directly downstream of the shikimate pathway targeted by glyphosate, but we also find that these alleles have pleiotropic roles in development and reproduction. We find particularly long haplotypes and low diversity associated with NTSR alleles, providing evidence for recent positive selection on this polygenic trait. Given that NTSR can explain 25% of the variance in phenotypic resistance, our findings imply that one can remain optimistic about the prospects for phenotypic prediction of resistance in weed populations, and that even for individuals harbouring multiple large-effect TSR mutations, NTSR may be of primary importance. Moreover, our findings suggest that selection from herbicides may have more widespread consequences on genomic diversity than initially assumed. Further work to functionally validate the candidate genes identified here will shed light on mechanism and consistency of these alleles in conferring resistance across populations.

## Data Accessibility

- DNA sequences available at European Nucleotide Archive (ENA) (project no. PRJEB31711); https://www.ebi.ac.uk/ena/browser/view/PRJEB31711
- Reference genome and its annotation available at CoGe (reference ID = 54057); https://genomevolution.org/coge/GenomeInfo.pl?gid=54057
- Scripts for data analysis available at https://github.com/jkreinz/NTSR-GWAS-MolEcol2020.
- Sampling locations and morphological data available in Sup Table 1.

## Author Contributions

JMK, SIW, and JRS designed the research; JMK conceived, analyzed, and wrote the paper; and JMK, SIW, JRS, PJT, and DW contributed feedback and edited the paper.

## Acknowledgements

We would like to thank Tyler Kent, Amardeep Singh, and George Sandler for helpful and fun discussions, Regina Baucom and two anonymous reviewers for thoughtful and useful feedback. JK was supported by a Society for the Study of Evolution Rosemary Grant and an NSERC PGS-D, JRS and SIW were supported by Discovery Grants from NSERC Canada, SIW was additionally supported by a Canada Research Chair in Population Genomics, and DW was supported by the Max Planck Society and Ministry of Science, Research and the Arts of Baden-Württemberg in the Regio-Research-Alliance “Yield Stability in Dynamic Environments”.

## Supplementary Figures

**Sup figure 1.**
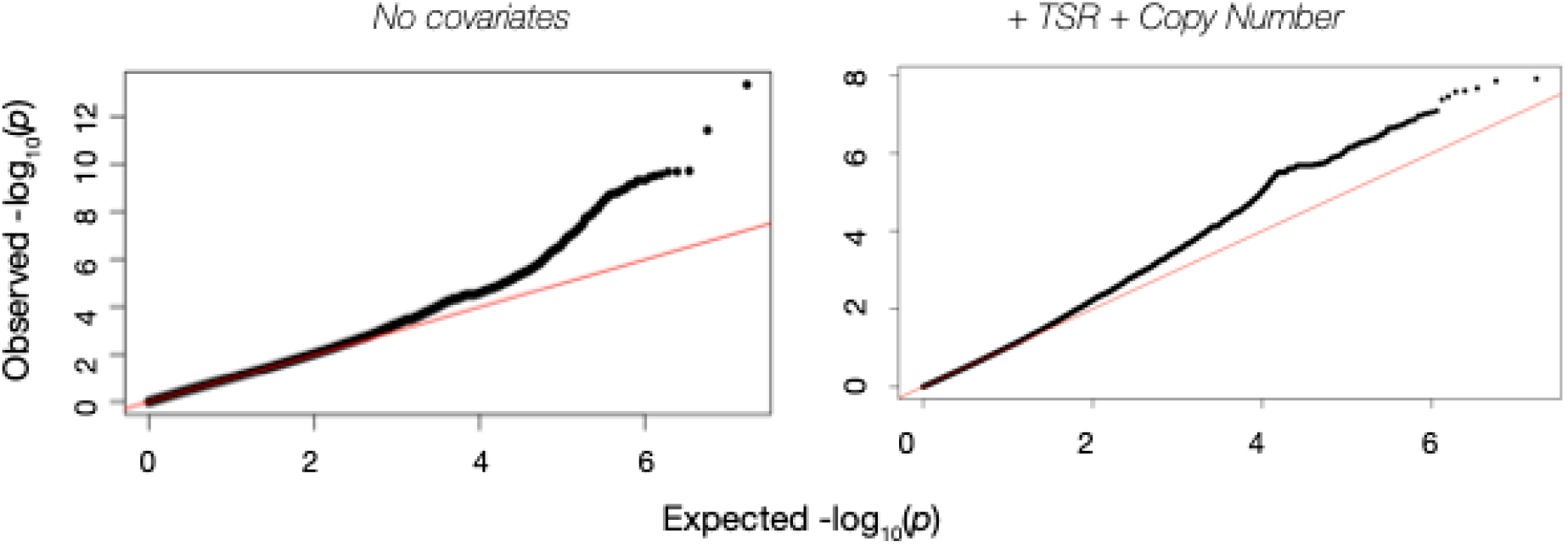
QQplots of observed versus expected p-values for GWA models of raw resistance levels (no covariates) and a model of non-target site resistance (using the EPSPS Pro106Ser substitution and EPSPS copy number as covariates).

**Sup Figure 2.**
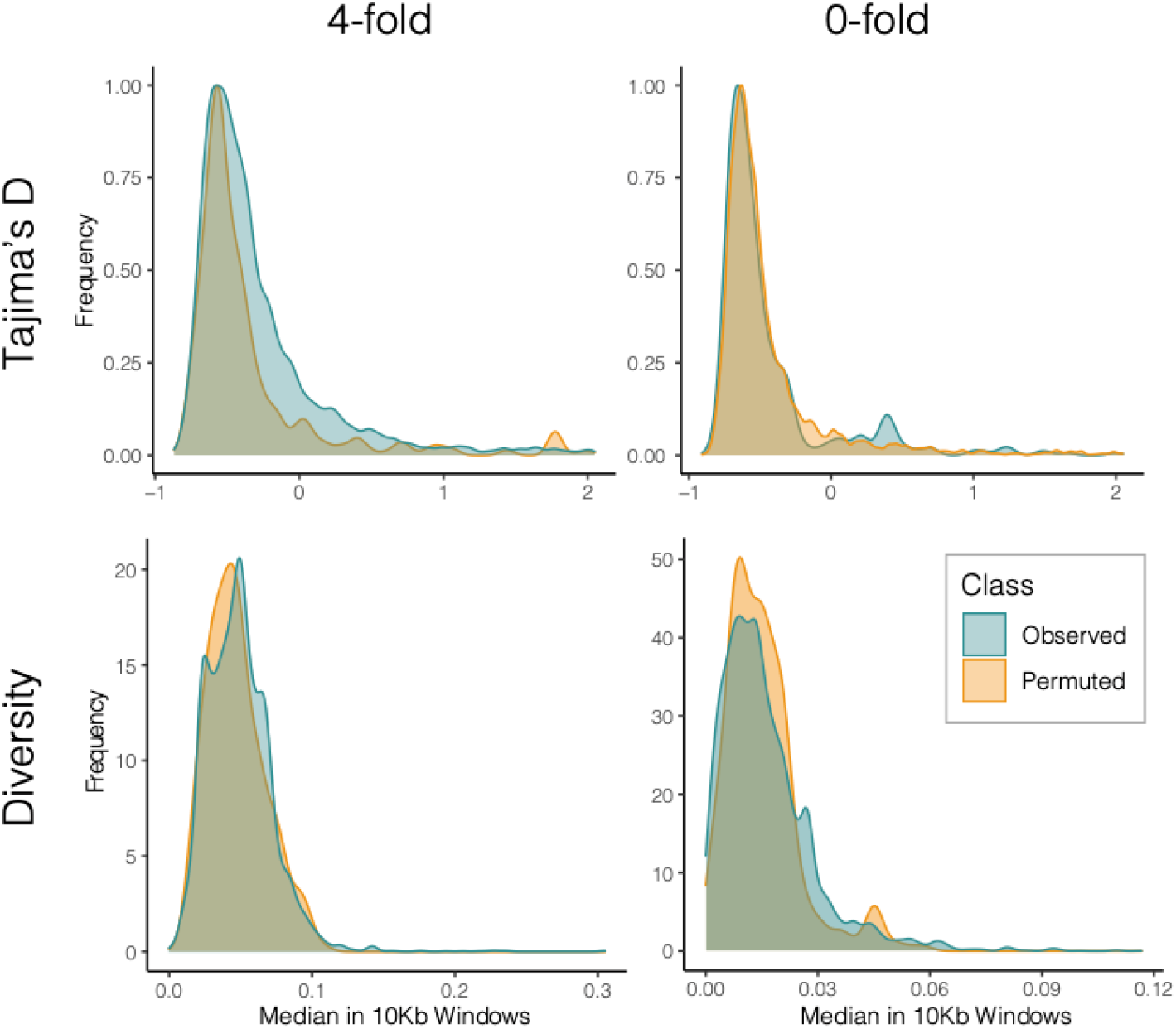
Distributions of 4 fold and 0 fold Tajima’s D and diversity in 10kb windows surrounding focal NTSR SNPs (across all scaffolds, or excluding the EPSPS-containing scaffold 5) relative to the permuted distributions.

